# Global State Changes Induce Non-reciprocal Reduction of Mutual Information in the Thalamocortical Network

**DOI:** 10.1101/213272

**Authors:** Ryan Verner, Matthew Banks, Edward Bartlett

## Abstract

Understanding the neural mechanisms of loss and recovery of consciousness remains an active area of scientific and clinical interest. Recent models and theories of consciousness implicate thalamus or neocortex as the dominant regions that control global state change, but experimentally testing these theories requires parsing the response properties of the complex thalamocortical network. In this work, we explore information processing using chronic recordings of multiunit activity between thalamus and cortex by targeting either the corticothalamic or thalamocortical pathway with direct electrical stimulation under subhypnotic and just-hypnotic states of isoflurane or dexmedetomidine induced unconsciousness. We found that mutual information between the stimulus and response in the corticothalamic circuit decreases with loss of consciousness, and that the change is not reciprocated in the thalamocortical direction. These changes in mutual information were moderately correlated with changes in evoked rate, and a strong effect of isoflurane on spontaneous and evoked rate accounts for a large portion of the change in mutual information. Further, we found cortical synchrony to increase under sub-and just-hypnotic doses of dexmedetomidine and isoflurane, suggesting cortical responses become more causally dependent on thalamocortical input at reduced levels of consciousness, or unconsciousness. We believe these changes suggest loss of consciousness is associated with a decrease in the number of available cortical network states, and may suggest cognitive unbinding from ascending sensory input.

## Introduction

Despite great efforts made toward understanding the molecular and neural mechanisms of anesthetic and sedative drugs (Antkowiak B 2001; Rudolph U and B Antkowiak 2004; Franks NP 2008), a unified theory of the network changes under various levels of consciousness or unconsciousness remains elusive. Changes in neural properties in the thalamocortical network are strongly implicated as a driving force for the loss and recovery of consciousness, and this network is central to theories regarding neural correlates of consciousness (Edelman GM 2003). However, the specific roles of cortex and thalamus in changing global brain states are unclear. The role of top-down versus bottom-up processing during loss-of-consciousness is of critical importance to understanding network changes in this circuit, as understanding the origin and mechanism of loss of consciousness in thalamus and neocortex may aid in the development of new intraoperative tools to monitor loss and regain of consciousness.

Previous work studying the neural basis for loss-of-consciousness has identified that anesthetics suppress activity in thalamus, leading to suppression of ascending input into sensory cortex (Ries CR and E Puil 1999; Alkire MT et al. 2000), and laid the groundwork for the *thalamic switch hypothesis* (Alkire MT et al. 2000). However, it has been demonstrated that suppression of cortical responses can be unrelated to awareness (Kerssens C et al. 2005), and in some cases, cortical sensory responses can be dramatically enhanced rather than suppressed during unconsciousness (Imas OA et al. 2005). In fact, anesthetics tend to have minimal effects on large scale organization and responsiveness of “core” thalamic nuclei that provide the bulk of ascending input into cortex (Jones EG 1998; Saalmann YB 2014), suggesting that thalamocortical signaling remains mostly intact during unconsciousness. While this ascending information continues to activate cortex, evidence suggests that it fails to activate an anesthetized cortical network with the same efficiency as a conscious one (Boly M et al. 2012; Liu X et al. 2012). The *information integration theory of consciousness* (Tononi G 2004) captures this premise, suggesting that integration of ascending information is necessary in order for a network to achieve consciousness. According to this theory, anesthetic drugs would need to work over a wide range of neural targets to reduce the number of available network states and connectivity between functional regions (Alkire MT et al. 2008). Thus, a closely related theory of anesthetic action, the *cognitive unbinding hypothesis* (Mashour GA 2013, 2013; Mashour GA and MT Alkire 2013), suggests that while ascending input arrives in cortical structures, corticocortical desynchrony prevents integration of information, creating lack of awareness through a disrupted perception of the environment. Both the information integration theory and cognitive unbinding hypothesis implicate changes or actions upon cortex are the primary drivers of loss-of-consciousness, suggesting that the observed thalamic suppression of ascending input is either a readout of anesthetic-induced unconsciousness, or possibly plays a complementary role. In either case, the stronger anesthetic effect on corticocortical connectivity (Massimini M et al. 2005; Ferrarelli F et al. 2010) and top-down pathways (Raz A et al. 2014) may be responsible for the modulation of cortical representation of sensory information.

We hypothesize that the thalamocortical and corticothalamic pathways will be differentially affected by anesthesia, and specifically at doses causing loss of consciousness. Information theoretic approaches in neural systems are a quick, high throughput mechanism to study information flow, and can be used with data from invasive (Nelken I et al. 1999; Nelken I et al. 2005; Chechik G et al. 2006) or non-invasive neural recordings (Jeong J et al. 2001; Schlögl A et al. 2002) in the operating room without the need for expensive scanning equipment or long time series. In the invasive preparation, neural spike trains can be studied as is, using the rate and temporal dynamics of a spike train to encode the stimulus. This is particularly important in the auditory thalamocortical system in which acoustic information is encoded in both spike rate and timing (Miller LM, MA Escabí, HL Read, et al. 2001; Miller LM, MA Escabí and CE Schreiner 2001; Miller LM et al. 2002; Bartlett EL and X Wang 2007; Wang X et al. 2008; Huetz C et al. 2009; Bartlett EL and X Wang 2011; Huetz C et al. 2011), so information theoretic approaches in this system should be designed to measure both of those response profiles.

While several studies have attempted to characterize information flow and cortical connectivity under various states of consciousness, most have used a form of peripheral, natural stimulation as a tool to activate the network. Under this experimental protocol, the stimulus passes through multiple subcortical nuclei and is already affected before reaching the thalamocortical circuit. Furthermore, feedback pathways cannot be directly assessed in the absence of feedforward pathways. Therefore in this work, we assess the information flow in the thalamocortical and corticothalamic directions using microstimulation of each nucleus, with simultaneous recording of the neural response to stimulation in the other nucleus. Using an information theoretic approach, we characterize the reduction in network information and changes in network synchrony which further validate the role of top-down pathways during loss-of-consciousness.

## Methods

Measurements were collected from 3 young adult male (age 4-6 months) Sprague-Dawley rats (Envigo, Indianapolis, IN). All procedures were approved by the Purdue Animal Care and Use Committee (PACUC 1204000631). The animals used in these studies were previously trained on an acoustic discrimination task that involved the discrimination of Gaussian noise from amplitude modulated Gaussian noise, but were naïve to the electrical stimuli used here. None of the rats had previous training related to tonal stimuli. All recordings were obtained under passive conditions and the rats were not performing any trained behaviors.

### Surgical Preparation

Aseptic animal surgeries were performed as previously described (Vetter RJ et al. 2004). Briefly, unconsciousness in rats was induced with isoflurane in air (5%) until loss of righting reflex was achieved, followed by delivery of a cocktail of ketamine (80 mg/kg) and dexmedetomidine (0.2 mg/kg) intraperitoneally to ensure a continued anesthetic depth of unconsciousness. Depth of anesthesia throughout surgery was monitored via frequent paw withdrawal reflex tests and maintained with additional intramuscular injections of ketamine upon reflex recovery. After anesthesia was confirmed, the animal was positioned in a stereotaxic head holder with hollow earbars to facilitate intraoperative electrophysiological recordings. Body temperature was maintained under anesthesia with a thermostatic water circulating pad and breathing and heart rate were monitored via pulse oximeter. After positioning in the stereotaxic head holder, the rat was shaved and the scalp was scrubbed for aseptic surgery.

An incision was made along the sagittal suture from bregma to lambda. The skin was loosened with blunt dissection and reflected to expose the dorsal face of the cranium. The right temporalis muscle was partially excised, with remaining portions reflected, to allow access to the cranium over the right auditory cortex. Four burr holes were drilled manually at positions rostral to bregma and caudal to lambda, all 3-4mm from midline, such that the bones screws would not interfere with the two electrode headstages. After the burr holes were complete, the holes were widened manually with a #56 hand drill to accommodate 0-80×1/8" stainless steel screws which would serve as the reference and ground for the electrode arrays. For a dorsal approach to the medial geniculate body, an approximately 2mm diameter craniotomy was centered at -5.5mm bregma, 3.5mm right of midline. The ventral division of the medial geniculate body was primarily targeted for this experiment using a linear 16-channel array of 703μm^2^ sites with 100μm site spacing (A1×16-Z16-10mm-100-703; NeuroNexus, Ann Arbor, MI), causing some of the sites (4-5) to reside in the dorsal division. These sites were never used for electrical stimulation, but recordings at these sites were analyzed along with sites from the ventral division. The 2mm diameter craniotomy over primary auditory cortex was performed caudally and dorsally to the squamosal suture. Primary auditory cortex was approached perpendicularly to the surface of the brain using a four shank electrode array with four, 703μm^2^ sites per shank (A4×4-HZ16-3mm-100-125-703_21mm; NeuroNexus), such that each shank resided along a different cortical column. The cortical electrode array was manually positioned to a depth of 1mm such that the most superficial site recorded putatively from layer 4. Surgical placement of the electrodes in both cortex and auditory thalamus were verified via intraoperative electrophysiological recordings of evoked responses to Gaussian noise bursts at 80dB SPL delivered through hollow earbars. After electrode placement was validated, the electrodes were locked in place with a two-part silicone elastomer (Kwik-Sil; World Precision Instruments) and the headcap was secured with UV curable dental acrylic. The animal was then allowed to fully recover from unconsciousness and was returned to his home cage. Post-operative assessments of weight and wound care were completed for 5 days after surgery, and daily administrations of meloxicam (1 mg/kg) in saline were delivered for postoperative pain management.

### Electrophysiological Recordings

Beginning three days after surgery, electrophysiological recordings were used to measure neural responses to acoustic and electrical microstimulation in both the medial geniculate body and primary auditory cortex. Recordings were collected from an acoustically and electrically isolated chamber (Industrial Acoustics Corporation). On each day of recording, the rat underwent a full bout of conditions of consciousness: wake, sub-hypnotic, just hypnotic, and recovery. For each drug condition, a loading period of 10 minutes was followed by an equilibration period. The next set of recordings only began after the animal had maintained a state of consciousness for a period of 10 minutes. One of two dosages of either isoflurane or dexmedetomidine induced altered states of consciousness while the rat was mildly restrained in a custom restraint apparatus (Supplemental Figure 1). Isoflurane (~0.6% for subhypnotic, ~0.9% for just hypnotic) was administered via vaporizer through a nosecone, while dexmedetomidine (0.08 mg/kg/hour 10 minute bolus + ~0.016mg/kg/hour dose for subhypnotic, 0.12 mg/kg/hour 10 minute bolus + ~0.024mg/kg/hour dose for just hypnotic) was delivered via tail vein catheterization, with only one drug being used on each recording day and isoflurane always being used on the day prior to dexmedetomidine. Approximately 20 hours passed between recording sessions. The animal’s state of consciousness was confirmed before and throughout recording sessions. Sub hypnotic levels of consciousness were confirmed with the presence of a paw reflex and righting reflex, while just hypnotic unconsciousness was verified by a maintained paw withdrawal reflex accompanied by loss of the righting reflex. In the case of dexmedetomidine, a reversal agent (Atipamezole, 1 mg/kg s.q.) was delivered to induce rapid recovery from the drug effect.

Signals from recording electrodes were sampled at ~25kHz on a zif digitizing headstage (Tucker-Davis Technologies) and passed to a RZ2 bioamp processor (Tucker-Davis Technologies) for filtering. Local field potentials were band-pass filtered on-line between 10Hz and 500Hz, while multiunit data were similarly filtered between 300Hz and 5000Hz. Trials which contained pronounced movement artifact were detected via cross correlation between channels and changes in RMS voltage, and these trials were not included in future analyses. Spikes were detected and sorted offline with waveclus (Quiroga RQ et al. 2004), however, insufficient single unit clusters were identified for processing. Thus, all data were analyzed as multiunit clusters. All of the electrodes used for this study underwent an activation protocol prior to implantation in order to deposit iridum oxide on their surfaces, thereby reducing impedance and increasing charge-carrying capacity (Robblee L et al. 1983). These features were necessary in order to stimulate on our electrodes while maintaining the integrity of the surrounding tissue and our electrode sites (Cogan SF et al. 2004; Merrill D et al. 2005), but the modification resulted in poorer unit isolation. Thus, in all recordings, multi-unit activity was analyzed.

### Electrical Stimulation

Electrical stimuli were sampled at ~25kHz and built in real time on a RX7 stimulator base station (Tucker-Davis Technologies) and delivered through an MS16 stimulus isolator (Tucker-Davis Technologies). Electrical pulses used for cortical stimulation composed of singular biphasic, symmetric, cathode-leading pulses of 205μs per phase in order to simulate corticothalamic feedback (Crandall SR et al. 2015). Electrical pulses used for thalamic stimulation were composed of a triplet of biphasic, symmetric, cathode-leading pulses of 205μs per phase at 300Hz, in order to simulate a thalamocortical burst volley (Swadlow HA and AG Gusev 2001). All electrical stimulation was performed on four sites in the cortical and thalamic arrays (Fig. 1A, gray circles). In the cortical arrays, the deepest sites on each array were selected such that the stimulated layers would be closest to cortical output layers. In the thalamic arrays, sites were selected with the intention of being equally spaced across the extent of the ventral division of the medial geniculate body to target the lemniscal auditory pathway, based on stereotaxic placement of the electrode array. This meant that the deepest site, along with three sites at 300μm intervals in the dorsal direction, were selected.

**Figure 1:**
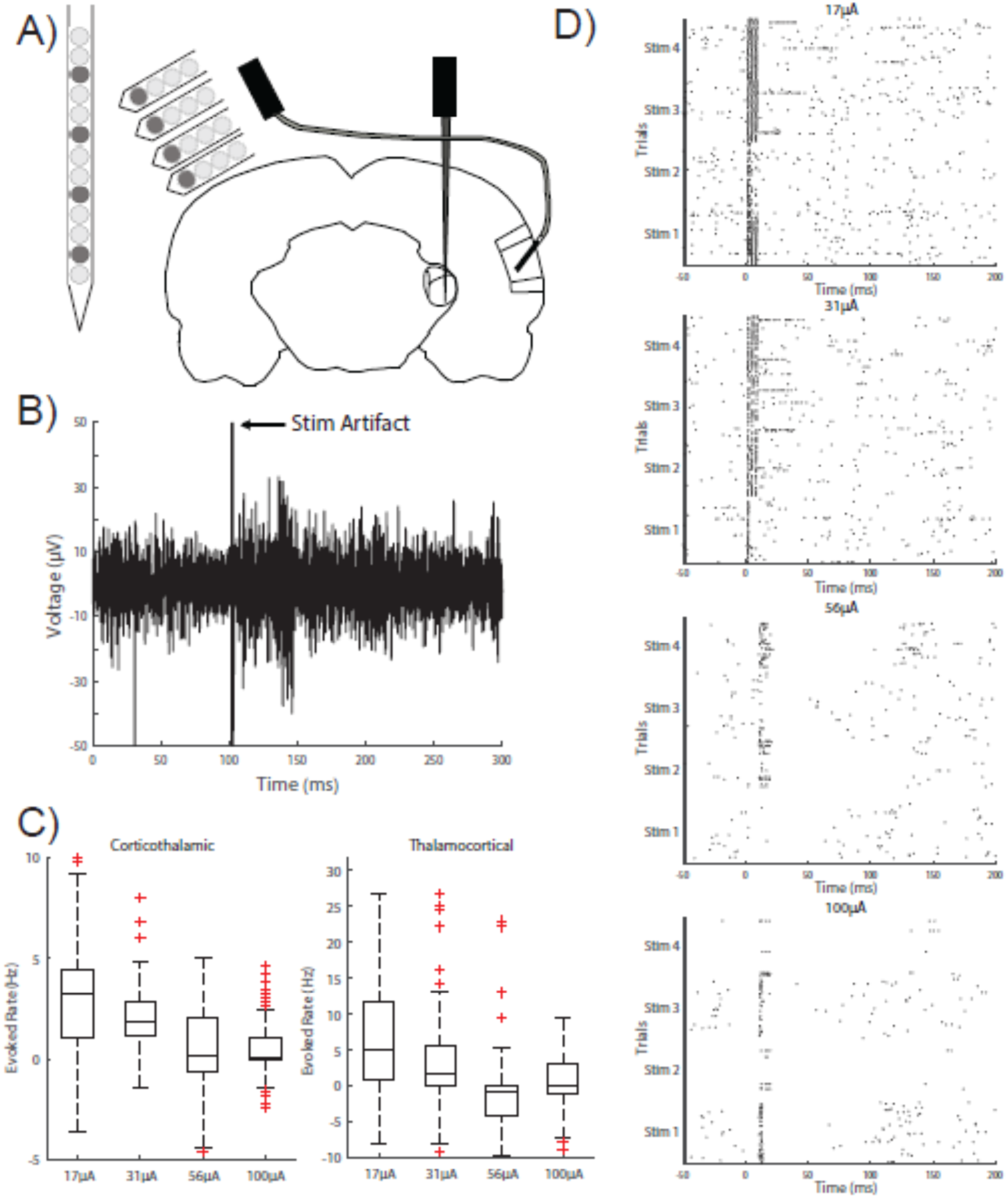
High intensity electrical stimulation invokes long-lasting suppression of responses. **A,** Electrodes were placed in primary auditory cortex and the ventral division of the medial geniculate body. Dark gray circles indicate stimulating electrodes (Stim 1-4, 1D), while responses were recorded on all electrodes from a given multichannel array. **B,** The stimulus artifact in our raw, multiunit activity data had excessively large amplitude compared to normal spiking data, and was easily detected and removed. **C,** Despite removal of the stimulus artifact, high intensity stimulation had a tendency to suppress responses to all stimulus channels. **D,** In extreme cases, such as at 56μA and 100μA, neural activity could be suppressed for over 100ms. For this reason, 31μA stimulation was used for future analysis.

### Information Theoretic Analysis

In order to study the raw spike trains, an incredibly large number of samples (~2^8^ per stimulus) is encouraged in order to reduce the upward bias and variability in information estimates (Treves A and S Panzeri 1995; Panzeri S et al. 2007). Binning responses can help to drive the variability of the information estimates down, even at a lower sample count, through response compression. When combined with a bias correction scheme, error in the information estimate is further diminished. A reliable estimate of mutual information between a stimulus and its evoked response allows for the measurement of small changes in mutual information right on the boundary of consciousness.

Information theoretic analysis was performed by taking 10 millisecond binned spike counts for the first 100ms after stimulus onset. For each of the four stimulus channels in either thalamus or cortex, binned responses in the opposite structure were collected in response to 50 repetitions. Each bin formed a letter of a response word. These responses were fed into a custom Matlab program, which calculated an estimate of the mutual information between the stimulus condition and the response profiles with the following equation:

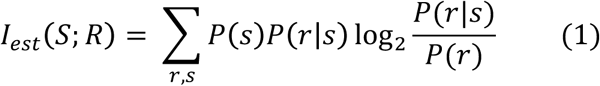

where S and R represent the stimuli and responses, respectively. To correct for bias in our MI estimate, the MI was calculated by removing the quadratic extrapolated bias (Treves A and S Panzeri 1995; Strong SP et al. 1998; Panzeri S *et al.* 2007) via the following equation:

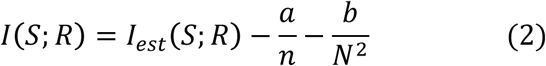

where *a* and *b* are parameters estimated by computing MI estimates (Equation 1) from fractions of the data, then fitting to a quadratic function of *1/N.* Such was the case for estimates of MI for each channel (Fig. 3). In some cases, channels lacked a pronounced response to stimuli and drove the bias-corrected MI to near zero. When a channel displayed consistently low evoked rates (<0.5 Hz) across all states of consciousness and stimulus intensities, the channel was believed to be unresponsive and was excluded from future analysis.

### Spike Time Tiling Coefficient

Mutual information analyses are typically used to assess the information conveyed by the spike timing in a neural code, but these measures do not directly assess the precision and reliability of spike timing within or across channels. During the course of information theoretic analysis, it was determined that changing spike rates during loss of consciousness had a major effect on mutual information. For this reason, spike time tiling coefficients were explored as an alternative method to examine pairwise correlations in spike trains (Cutts CS and SJ Eglen 2014). Spike time tiling coefficients were determined for each of the recording channels by comparing responses to stimulation on sequential trials using the following equation:

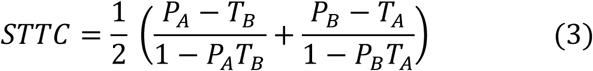

where *P*_*A*_ and *P*_*B*_ represent spike probabilities, and *T*_*A*_ and *T*_*B*_ represent the fraction of tiled time analyzed per trial. For example, if trial *A* were being compared to trial B, spikes were first detected in *A* and *B* and a tiling window is set around each spike with a width of *2*Δ*t* (in this work, Δ*t*=5ms). Using the duration of the trial, and the total duration of tiled time, *T*_*A*_ and *T*_*B*_ were calculated. Next, *P*_*A*_ was calculated by assessing the proportion of spikes in trial *A* that occur during the tiled time of trial B. The same was done for *P*_*B*_. This process was repeated for every set of trials for a given channel and the result is averaged for that channel.

### Statistical Analysis

Statistical analyses were performed using the MATLAB Statistics Toolbox using linear mixed effects models. Initial exploration of the data revealed a significant intrasubject variance which was accounted for as a random effect in our model:

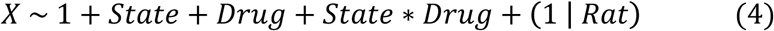

Where X was the dependent variable: mutual information or spike time tiling coefficient. Beyond the linear mixed effects models, regression analyses were performed to identify relationships between dependent variables, such as evoked rate and mutual information.

## Results

In our experiment, data were collected in response to a range of stimulation intensities. The range (17μA to 100μA) was initially selected based on behavioral thresholds of perception of electrical stimulation collected from other rats (data not shown), with the intention of collecting some neural responses to sub-threshold stimulation and supra-threshold stimulation. In practice, the spike rasters revealed a suprathreshold neural response to all levels of stimulation, indicating that behavioral salience of electrical stimulation in our task requires greater stimulation intensity than low-level, focal activation of neural circuitry (Fig. 1D). In fact, the neural responses to 100uA stimulation, and often to 56uA stimulation, revealed prolonged inhibitory effects which we believed to be linked to wideband forward suppression (Fig. 1D). Further evidence of this effect can be seen in the average evoked rates in response to increasing levels of stimulation, where the responses to higher stimulus intensities were driven closer to zero, sometimes below spontaneous rate, in both the thalamocortical and corticothalamic directions (Fig. 1C). We present data collected at the 31μA intensity for the remainder of the article, as it represented the highest stimulation intensity than did not evoke a strong inhibitory effect.

### Changes in Rate with Drug and State

In our study, we elected to use multiple anesthetic or sedative drugs in order to identify common changes in information transfer between each drug condition. We found dexmedetomidine to induce a reduction of spontaneous rate in the just hypnotic condition, while isoflurane induced strong reductions of spontaneous rate in both drug levels (Fig. 2). These changes in spontaneous rate occurred in both the corticothalamic and thalamocortical directions. Dexmedetomidine didn’t not exhibit a strong effect on evoked rate, and isoflurane’s effect on evoked rate was only observed in the corticothalamic direction (Fig. 2). While the changes in spontaneous rate are supported by previous work, there is not a strong precedent in the literature for drug effects on firing rates evoked by direct activation of these network pathways.

**Figure 2:**
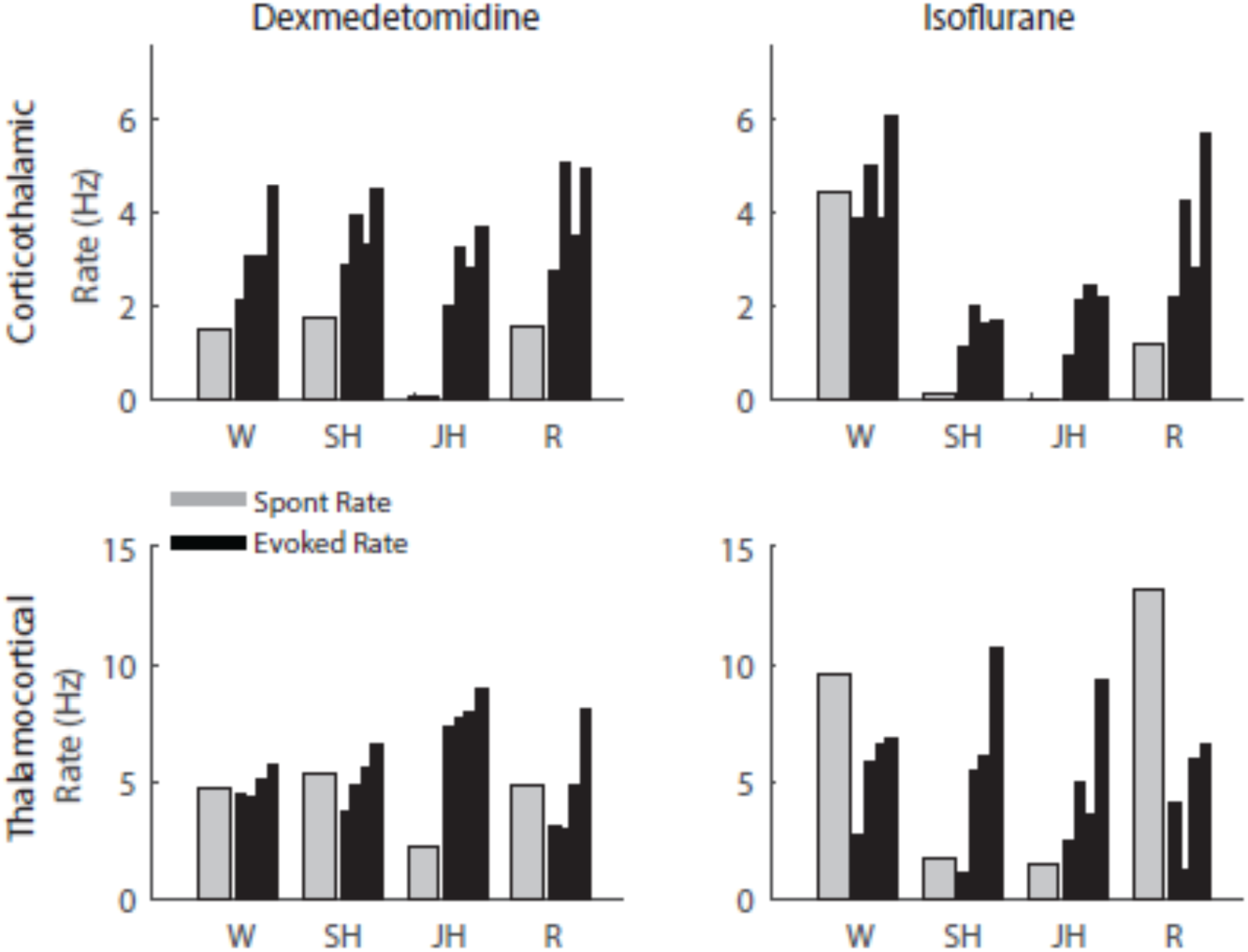
Isoflurane causes a characteristic reduction of spontaneous rate. Dexmedetomidine and Isoflurane were used in comparison in order to control for the known effects of Isoflurane on firing rates. Bars represent spontaneous (gray) or evoked (black) rates, where each black bar represents a rate response to a specifici stimulation channel. In both corticothalamic and thalamocortical directions, Isoflurane had a marked effect on rate at the sub- and just-hypnotic doses. These effects were not as prevalent in Dexmedetomidine-induced unconsciousness.

### Channel-specific Mutual Information

In order to assess channel-specific changes in mutual information, mutual information was calculated on each channel at each state of consciousness by analyzing the neural response to each of four stimulation channels. In figure 3, one channel per drug condition and network direction (each column) was selected to display some of the typical response profiles and how those profiles changed with brain state. In most cases, wake and recovery conditions exhibited the highest spontaneous rates and sometimes had stronger sustained response profiles to stimulation. In recordings of the sub-and just hypnotic levels of each drugs, responses either became more onset-driven or disappeared (Fig. 3).

**Figure 3:**
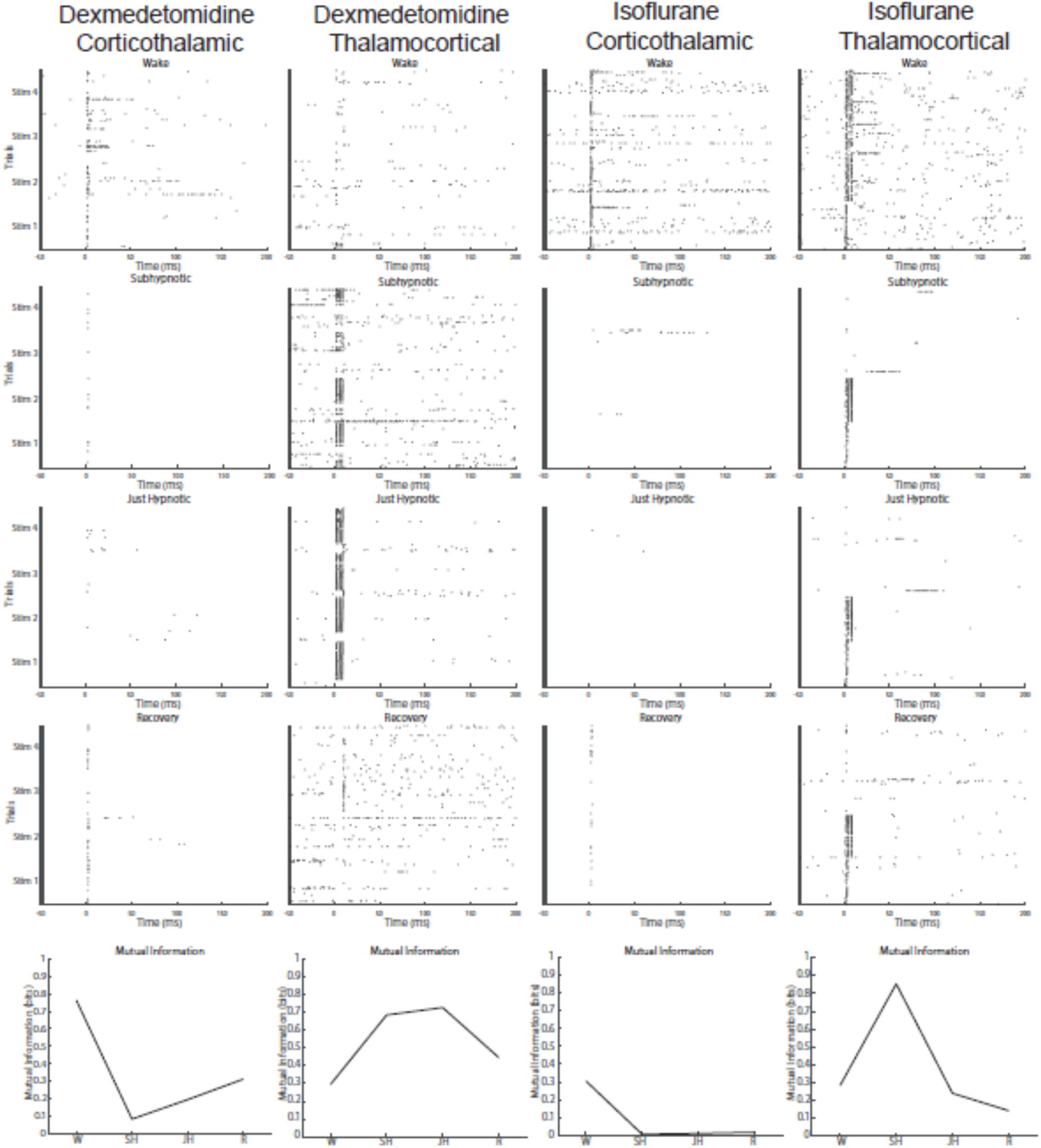
Calculation of channel-specific Mutual Information. For each channel, spike probabilities per bin were calculated for the first 100 milliseconds after stimulus onset. At each state of consciousness, for each channel, a mutual information was calculated. Generally, responses occurred in the first 2 bins (first 20ms), but in the wake and recovery conditions, responses were more distributed.

### Non-reciprocal Reduction of Corticothalamic Mutual Information

While many groups have assessed mutual information in auditory systems (Nelken I *et al.* 2005; Chechik G *et al.* 2006; Atencio CA et al. 2008; Gaucher Q et al. 2013), a large majority of the work is restricted to measuring the neural responses to natural stimuli. In our study, we sought to characterize changes in mutual information along feedforward and feedback pathways directly.

Our results show a non-reciprocal reduction of mutual information in the corticothalamic pathway, which was most significant under isoflurane-induced unconsciousness (Fig. 4). While the dexmedetomidine corticothalamic data was not found to be significant, the difference between the wake and just hypnotic level of unconsciousness approached significance (p=.07), and, if analyzed as a continuous trend from wake to unconsciousness (just hypnotic), the effect was found to be significant (p<.05). Further, we found a significant difference in how isoflurane modulated this mutual information change as compared to dexmedetomidine in both the corticothalamic and thalamocortical directions (p<.001). Due to the bin sizes used to calculate mutual information, we were concerned our metric might be highly sensitive to rate changes. Thus, we analyzed the change in mutual information versus the change in rate under each drug condition, comparing the sub-and just-hypnotic states to the wake state. We found a correlation between mutual information and rate changes (Fig. 5). Rate does drive the majority of the mutual information effect in most cases. However, the r^2^ values indicate that at least 40% of the variance in these mutual information changes cannot be attributed to changes in rate.

**Figure 4:**
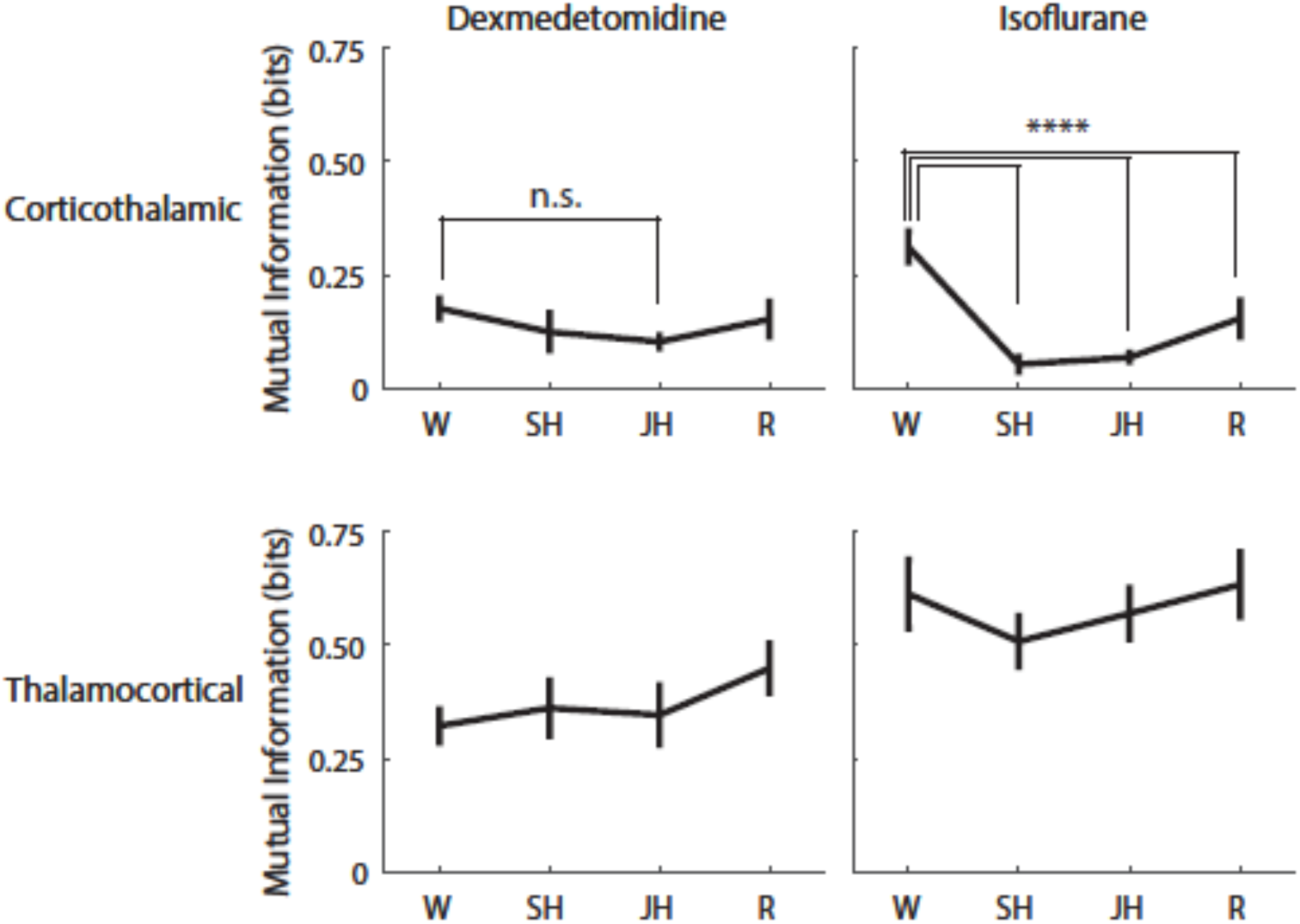
Corticothalamic Mutual Information decreases with loss of consciousness

**Figure 5:**
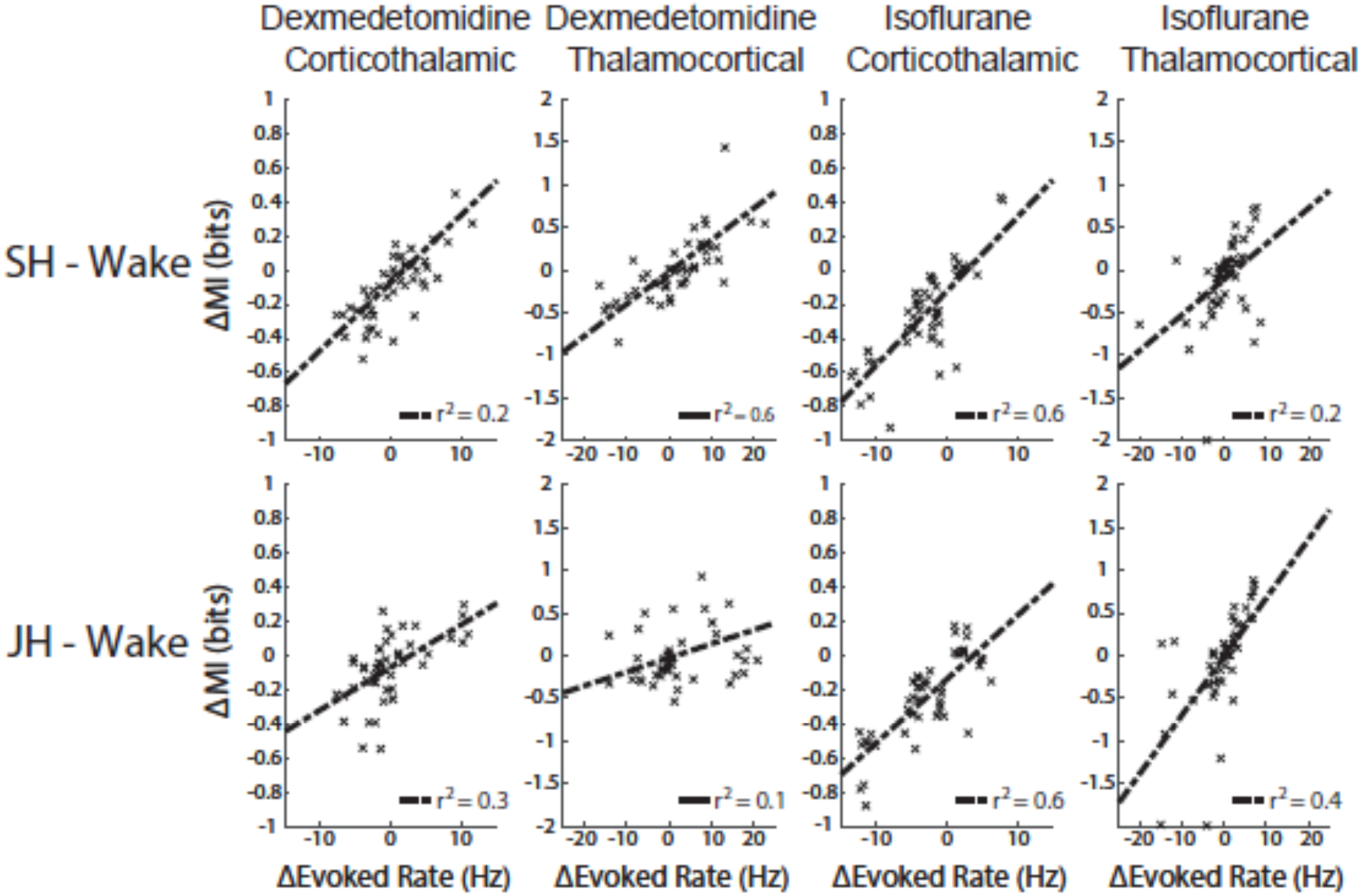
Changes in mutual information correlate with changes in rate

We also note higher overall thalamocortical information than corticothalamic information at all states of consciousness. Due to our bin size and the differences in stimulation profiles (thalamocortical burst vs. single corticothalamic pulse), it is likely that cortical responses to the thalamocortical stimulation existed in the first two bins, rather than just the first bin as would be the case in thalamic responses. This is the likely cause of the difference in overall magnitude. Because of inherent experimental differences in stimulation profiles, we only examine differences between drug conditions and dosage levels for a given pathway, rather than comparing the pathways directly.

### Pairwise Dependencies of Cortical Responses Increase with Loss of Consciousness

Due to the correlation between rate and mutual information, a rate-insensitive method to detect pairwise correlations and synchrony was selected to analyze the responses further. We found cortical spike time tiling coefficients to increase at the sub-and just-hypnotic levels of unconsciousness under both drugs (Fig. 6), however, this trend was not different between the two anesthetic drugs as it was using the mutual information metric. In auditory thalamus, the effects were far more variable and did not follow the same trend we saw in cortex, so it is possible that factors other than unconsciousness or anesthetic load are influencing these changes in auditory thalamus. Generally speaking, wakeful responses have lower spike time tiling coefficients than anesthetized or drowsy responses, and it appears as though cortical circuits return to the wakeful response profile more readily than thalamic circuits.

**Figure 6:**
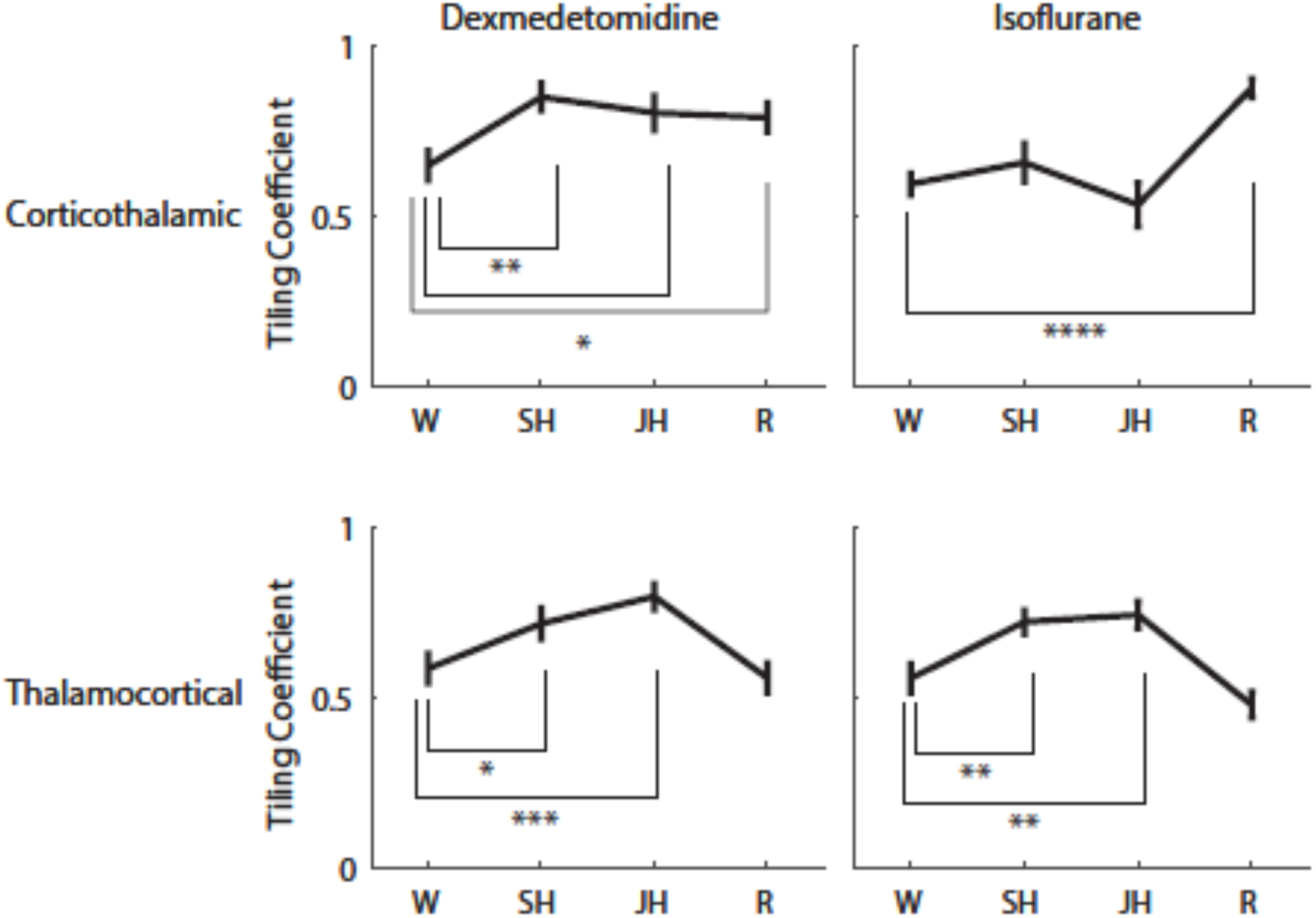
Cortical Spike Time Tiling Coefficient increases with loss of consciousness

### Network Response Synchrony Decorrelated From Mutual Information

We found changes in mutual information and spike time tiling coefficients to be uncorrelated in our data (Fig. 7). As expected, spike time tiling coefficients were generally found to be decorrelated from rate (not shown), and given the correlation between rate and mutual information, it is unsurprising that spike time tiling coefficient and mutual information would be uncorrelated in our data.

**Figure 7:**
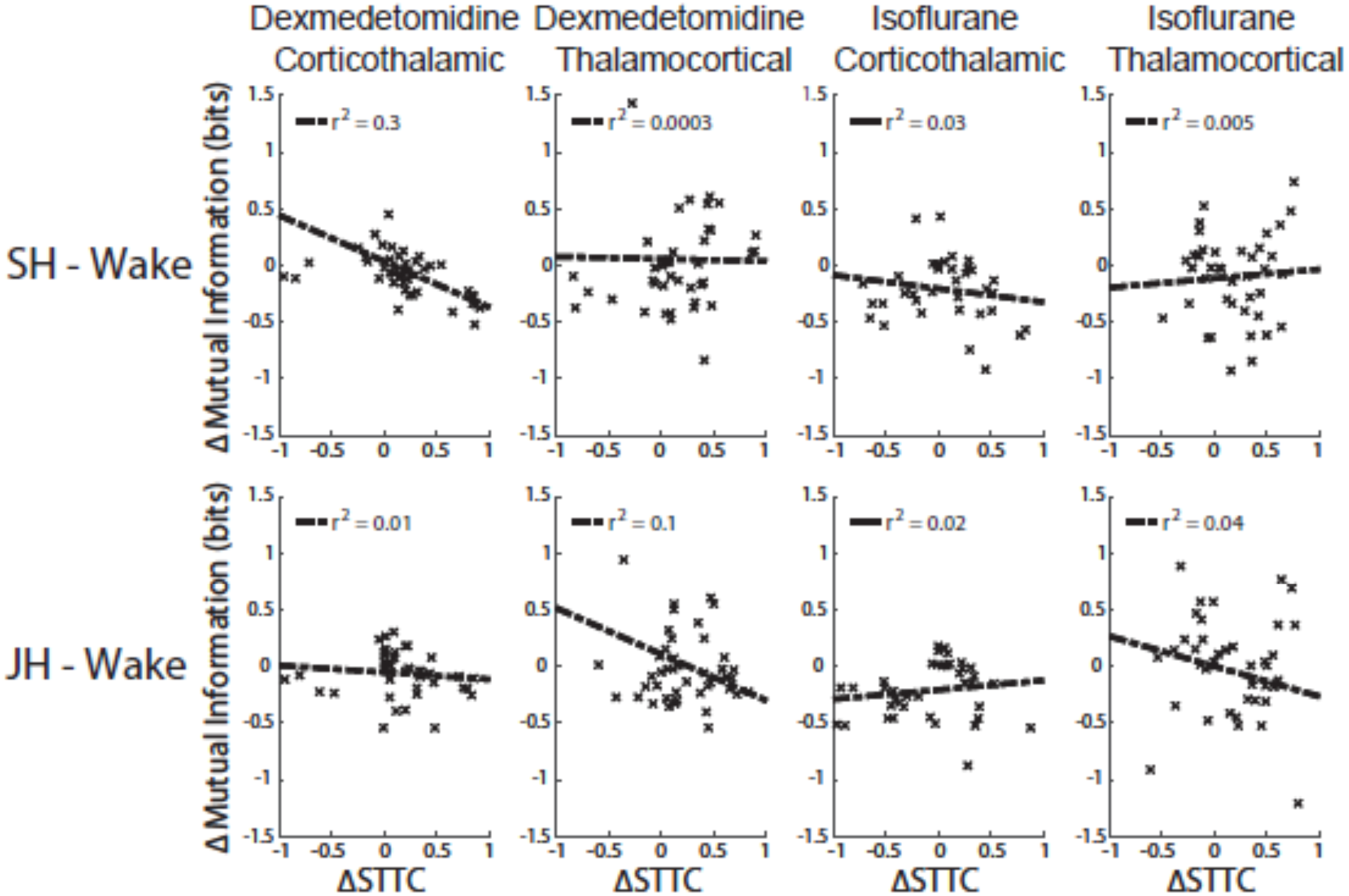
Changes in mutual information and spike time tiling coefficient are not correlated.

## Discussion

This work represents a focused approach to assessing information flow and changes in network properties during loss of consciousness. To this end, we electrically activated one nucleus of the thalamocortical circuit and recorded the resultant effects on the other end of the circuit. Unlike typical studies which assess information between two anatomically connected neurons or networks by comparing two spike trains, we could directly manipulate the pathway in a way that allowed us to approach an deterministic experimental approximation of “effective information” (Tononi G 2004). This method also allowed us to assess changes in network synchrony in response to brief, “wideband” stimulation – a measure of causal dependency to input streams.

We employed anesthetic drugs to achieve global brain state changes as a surrogate for natural loss of consciousness in a timely and repeatable manner. These drugs have known effects on neural circuitry at the single neuron and network levels. We chose to use two drugs, the volatile anesthetic isoflurane and the sedative dexmedetomidine, such that we could examine the effects of changes in unconsciousness that transcend specific pharmacokinetics of each drug.

### Drug-induced Changes in Firing Rate

The general anesthetic isoflurane, which has diverse effects on GABA, glutamate, and glycine receptors (Antkowiak B 2001; Grasshoff C and B Antkowiak 2006; Franks NP 2008; Shelton KL and KL Nicholson 2010; Ogawa SK et al. 2011), is well-known for decreasing spontaneous and evoked firing rates at in neocortex (Hentschke H et al. 2005; Land R et al. 2012; Sitdikova G et al. 2014; Noda T and H Takahashi 2015). In our data, we see spontaneous and evoked rate reductions under sub-and just-hypnotic doses of isoflurane that are consistent with the body of previous work (Fig. 2). Changes in rate under isoflurane are representative of a generally inhibitory effect of the drug on neural activity. By contrast, the α2-adrenergic receptor agonist dexmedetomidine has been shown to have a much lesser effect on evoked firing rates in the auditory system (Ter-Mikaelian M et al. 2007). In our data, we see a small decrease in spontaneous rate at the sub-hypnotic dose of dexmedetomidine, and a far greater decrease at the just-hypnotic dose (Fig. 2). These changes do not appear in the evoked firing rate data, which is consistent with documented effects of the drug.

### Change and MI and Change in Rate

Depending on how it is calculated, mutual information can be quite sensitive to changes in total firing rate. In our case, mutual information is computed by averaging spike count per 10 millisecond bin, and as such, higher firing rates may result in higher response entropy, and thus higher total information. Dexmedetomidine was selected to compare with isoflurane specifically due to differences of effect on firing rate in order to create a control for these known effects of rate on mutual information. In our data, we can assign approximately 60% of the changes in mutual information to evoked rate changes under loss of consciousness (Fig. 5), and in both the corticothalamic and thalamocortical pathways, we identified a significant effect of the drug used on resultant MI (Fig. 4). In the corticothalamic pathway, the interaction effect of Isoflurane on loss of consciousness was also significant (p<.001). We believe this interaction effect is due to the strong inverse correlation between Isoflurane concentration and firing rate, which we document in our findings (Fig. 2). However, the remaining 40% or more of variance associated with mutual information changes, as well as the near-significant categorical trend of mutual information reduction in dexmedetomidine-induced loss of consciousness, suggests the non-reciprocal disruption of information flow is associated with loss of consciousness and is not specifically an effect of certain anesthetic drugs.

### Validation of Competing Theories of Consciousness

We hypothesized that the thalamocortical and corticothalamic pathways will be differentially affected by anesthesia, and specifically at doses causing loss of consciousness. Our previous work is consistent with both cognitive unbinding and loss of information integration as primary drivers of loss of consciousness, especially in isoflurane-induced unconsciousness (Raz A *et al.* 2014).

Our data reveal an increase in cortical synchrony, as well as pairwise spike dependencies during input volleys, in sub-and just-hypnotic concentrations of anesthetics drugs (Fig. 6). This change in cortical responses is indicative of a reduction of information integration, as sustained cortical processing of stimuli happens in networks distributed throughout a cortical column over several hundred milliseconds (Wehr M and AM Zador 2005; Sadagopan S and X Wang 2010). Further, these changes indicate that thalamic depression is an unlikely initiator of loss of consciousness, as thalamocortical information transmission was unchanged at the low doses of sedatives and anesthetics we used (Fig. 4). We should reiterate, however, that thalamus could (and likely does) play a role in loss of consciousness. In fact, it is possible that poorly informative feedback from cortex drives stronger, wideband feedforward suppression, disrupting normal feed-forward processing and driving the entire thalamocortical network deeper into unconsciousness.

### Study Limitations

While our work provides specific insight into the roles that ascending and descending pathways play in drug-induced loss of consciousness, using electrical stimulation to activate these pathways introduced limitations to our study. Most notably, the stimulation artifact corrupted the local field potentials for tens of milliseconds after stimulus offset. This ringing artifact was due to our current-controlled stimulus with exceptionally sharp offset passing through our low-pass hardware filter. The only viable frequencies for local field potential analyses were approximately 15 Hz and above, and even at those frequencies, spectral analyses near the stimulus window were still heavily corrupted by the stimulus artifact. Thus, despite our initial interest in providing comparative results between local field and multiunit responses, our work was necessarily limited to the multiunit analysis.

In terms of the results that we were able to provide, however, the duration of each day’s experiment was a critical limitation. As noted earlier, gathering a sufficient number of responses to each stimulus condition is critically important for mitigating bias in mutual information estimation. In this work, our desire to collect responses at multiple stimulation intensities, at multiple drug levels, in both network directions, caused experiments to run for exceedingly long periods of time even with non-ideal numbers of repetitions per stimulus. We recommend that future examination of this effect focus on a smaller number of independent variables, in order to reduce experiment duration and information bias.

